# The Camouflage Machine: Optimising protective colouration using deep learning with genetic algorithms

**DOI:** 10.1101/2020.01.12.903484

**Authors:** J. G. Fennell, L. Talas, R. J. Baddeley, I. C. Cuthill, N. E. Scott-Samuel

## Abstract

The essential problem in visual detection is separating an object from its background. Whether in nature or human conflict, camouflage aims to make the problem harder, while conspicuous signals (e.g. for warning or mate attraction) require the opposite. Our goal is to provide a reliable method for identifying the hardest and easiest to find patterns, for any given environment. The problem is challenging because the parameter space provided by varying natural scenes and potential patterns is vast. Here we successfully solve the problem using deep learning with genetic algorithms and illustrate our solution by identifying appropriate patterns in two environments. To show the generality of our approach, we do so for both trichromatic and dichromatic visual systems. Patterns were validated using human participants; those identified as the best camouflage were significantly harder to find than a widely adopted military camouflage pattern, while those identified as most conspicuous were significantly easier than other patterns. Our method, dubbed the ‘Camouflage Machine’, will be a useful tool for those interested in identifying the most effective patterns in a given context.

## Introduction

Colouration has been an exceptionally useful phenotype for biological research, from genetics and development to ecology and evolution^1^. However, moving beyond colour per se to patterning, it is also a difficult phenotype to characterise. While a colour can be represented in a relatively low-dimensional space based on spectral characteristics, photoreceptor sensitivities or psychophysical measurements^2^, a pattern (a visual texture) is a high-dimensional attribute^3,4^. The problem of characterisation is particularly acute when the interest is in a colour pattern shaped by the perception of signal receivers as well as ecology, as will be the case for camouflage or biological signals, because a single colour pattern may need to be represented in multiple perceptual spaces. Yet characterisation is just the starting point for an even greater problem if we wish to evaluate the effectiveness of a colour pattern for a visual function such as camouflage or signalling: the difficulty of searching a high-dimensional space for the optimal solution. Here, we show how residual deep neural networks (RDNNs)^5^, combined with genetic algorithms, can be harnessed to classical psychophysical techniques to search a high-dimensional spatiochromatic space for optimal camouflage and signalling patterns.

To help understand the depth of the problem, take the study of animal camouflage. Research has typically experimentally tested a small set of pattern types relevant to a specific functional hypothesis, or has identified ecological correlates of extant patterns, i.e. patterns seen in nature^6,7,8,9^. Such studies necessarily omit possible patterns that evolution has not realised because of phylogenetic or developmental constraints, and so cannot identify the influence (if any) of such constraints. Furthermore, without comparison to the optimal pattern(s), it is hard to identify the extent to which an observed pattern is subject to trade-offs with other functions such as thermoregulation or UV-protection^10,1^. Defining a framework that could effectively characterise patterns, and realistically evaluate these patterns in terms of their visibility in a given context, would be an exceptionally useful research tool. Not only would this provide insight into animal camouflage, but also allow the assessment of whether the signals that animals use to display, variously, their qualities to mates or unprofitability to predators, are optimised for conspicuity. These may be subject to trade-offs that render maximal conspicuity suboptimal and/or favour tuning of the signal to particular receivers at particular distances^11,12,13,14^. In the human domain, our method may be useful in the development of bespoke camouflage for specific contexts, maximizing the visibility of warning signs, or helping to reduce visual clutter due to infrastructure.

The main purpose of this paper is to propose and test a new method that can identify effective patterns for a given environment. Depending on context and requirements, the method is applicable for finding patterns that will be effective either for camouflage or to be highly conspicuous. Historically, methods used to evaluate patterns tend to be based on binary comparison or measuring detection speed and accuracy, typically on computer screens. This is useful if there are only a few patterns to compare, but if the aim is not to constrain the space of possible patterns artificially then this approach is inadequate. Our method proposes gathering data, using human participants, on a subset of the parameter space and then, using residual deep neural networks^5^ to interpolate between samples, to predict the difficulty in finding empirically untested patterns.

To make the method highly applicable to real-world scenarios, we constructed naturalistic stimuli and, for realism, projected them on a screen large enough to fill the visual field. We used backgrounds taken from photographs of both temperate forest and scrub desert with foreground occlusion layers and targets inserted into the scenes using blue screening (“chroma key”), a method commonly employed in the film industry. We were also keen that the textures on the targets that we used had biological plausibility. To achieve this, we used two-component reaction-diffusion equations. These systems, originally proposed by Alan Turing^15,16^, consist of semi-linear parabolic partial differential equations capable of creating a vast array of textures, including the camouflage patterns of animals^17,18^. Textures were colour-mapped using one colour (represented as an RGB triplet) for each of the two components, creating two-colour, natural-looking patterns. Besides the human visual system, which is trichromatic, we also tested simulated dichromatic stimuli, because most mammals are dichromats^19^.

Targets in our main experiments were constructed using nine dimensions (three for each of the two colours and three for texture), resulting in a parameter space containing a total of 6.18 × 10^17^ possible patterns. Since our parameter space was so large, we were unable to exhaustively or randomly select targets with sufficient diversity. Therefore, we implemented a genetic algorithm (GA) to optimise the colour and texture parameters, based on participants’ responses trial by trial, for hardest or easiest to see stimuli^20^. Our first three experiments were pilot experiments conducted to validate the genetic algorithm using an increasing number of optimised dimensions: the first experiment optimised for targets with single trichromatic colours; the next experiment tested the optimiser with greyscale reaction-diffusion textures; and the final pilot optimised for two colours, but using a fixed pattern. Our hypothesis was that, over experimental generations, the reaction times to targets would gradually increase or decrease depending whether targets were optimised for camouflage or conspicuity, respectively. Analysis using General Linear Mixed Models showed support for a working GA and we proceeded with our main experiment.

The main experiment followed a 2×2 design with two types of backgrounds (temperate forest or semi-arid desert) and two colour vision conditions (trichromatic or simulated dichromatic); examples of the stimuli are illustrated in Figure 1. The results of the main experiment were used to train RDNNs which we then used to predict reaction times (a measure of difficulty) for a far greater number of patterns than had been observed by the human participants. A final experiment was conducted to validate the method.

**Figure 1.**
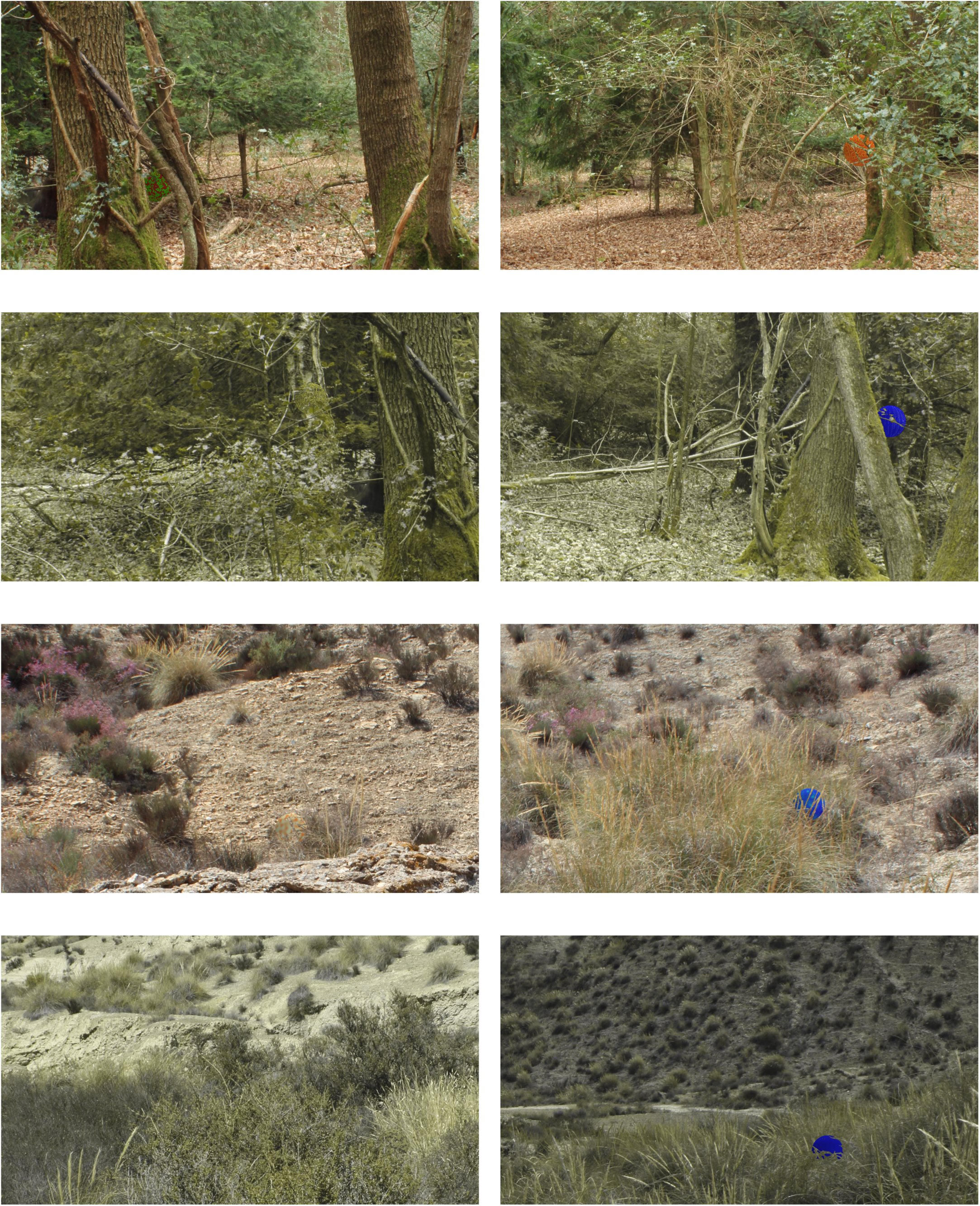
Examples of experimental stimuli shown in experiments 4a-d. From top to bottom each row depicts the following conditions: trichromatic temperate forest; dichromatic temperate forest; trichromatic semi-arid desert; dichromatic semi-arid desert. Columns illustrate examples of hard (left) and easy to see (right) targets.

## Results

### Genetic algorithm optimises for camouflage or conspicuity

The three pilot experiments confirmed that the genetic algorithm was capable of optimising target colour and texture for both concealment and high visibility. General linear mixed models (GLMMs) showed that trials became significantly harder over time when optimising for concealment, while optimising for visibility yielded easier to find targets (Table 1). The effects of trial number on log-transformed reaction times were analysed by fitting general linear mixed models using the lme4 package^21^ in R^22^. Nested models were compared using the change in deviance on removal of a term. A positive estimate coupled with a significant p-value suggested that targets became harder to see over the course of the experiment, while negative estimates indicated that targets became easier to see. It should be noted that estimates and standard deviations are presented as log-transformed reaction times. For example, an estimate of 1.86e-4 indicates that the target in the final trial was approximately one second harder to find than the target in the initial trial. In the main experiment, the optimiser produced significantly harder/easier results according to settings (see Table 1, experiments 4a-d), except in the dichromat desert condition optimised for easiest to see targets (p = 0.5321); we address this in the discussion below.

**Table 1.** Details of the GLMM analysis to determine whether the GA is effective, for all experiments. Experiments 1-3 are pilot experiments and experiments 4a-d are the main experiment.

| Experiment | Colour | Optimised for | N | ΔDeviance | Df | p | Estimate | Std |
| --- | --- | --- | --- | --- | --- | --- | --- | --- |
| Experiment 1 | Trichromat | Hardest | 5 | 17.293 | 1 | 3.20e-05 | 1.86E-04 | 4.46E-05 |
|  |  | Easiest | 5 | 22.510 | 1 | 2.09e-06 | -1.38E-04 | 2.89E-05 |
| Experiment 2 | Monochrome | Hardest | 5 | 11.552 | 1 | 6.77e-04 | 3.17E-04 | 9.32E-05 |
|  |  | Easiest | 5 | 317.050 | 1 | < 2.20e-16 | -1.14E-03 | 6.21E-05 |
| Experiment 3 | Trichromat | Hardest | 10 | 101.520 | 1 | < 2.2e-16 | 1.47E-04 | 1.46E-05 |
| Experiment 4a | Trichromat | Hardest | 5 | 10.771 | 1 | 0.001031 | 6.53E-05 | 1.99E-05 |
|  |  | Easiest | 5 | 16.633 | 1 | 4.54e-05 | -5.34E-05 | 1.31E-05 |
| Experiment 4b | Dichromat | Hardest | 5 | 47.902 | 1 | 4.48e-12 | 2.10E-04 | 3.03E-05 |
|  |  | Easiest | 5 | 16.612 | 1 | 4.59e-05 | -6.34E-05 | 1.56E-05 |
| Experiment 4c | Trichromat | Hardest | 5 | 7.565 | 1 | 0.005951 | 4.72E-05 | 1.71E-05 |
|  |  | Easiest | 5 | 156.360 | 1 | < 2.2e-16 | -1.59E-04 | 1.26E-05 |
| Experiment 4d | Dichromat | Hardest | 5 | 5.160 | 1 | 0.02317 | 4.83E-05 | 2.13E-05 |
|  |  | Easiest | 5 | 0.390 | 1 | 0.5321 | -9.02E-06 | 1.44E-05 |

### Residual deep neural networks were built to model the full parameter space

RDNNs were implemented in Keras 2.1.2^23^ utilising the neural network library TensorFlow 1.5.0^5^ and were trained with all of the samples collected from the main experiment. To provide for a measure of accuracy in our predictions (an estimate of standard error) we created 100 bootstraps of our networks for each of the four conditions. The bootstrap method is a test or metric that uses random sampling with replacement. The bootstrap method allows assignment of accuracy measures, defined here in terms of variance and is particularly useful when the value of interest is, as in the present case, a complicated function^24^. By averaging the bootstrapped networks predictions we calculate both a data-dependent smoothing of the reaction time function and an estimate of our certainty of its estimate. Each network was trained on a random sample of 90% of the data and validated with the remaining 10%.

### Artificial observers were created to predict pattern difficulty

Predicting the full parameter space poses a computational challenge due the vastness of the space. We therefore created 100 “artificial observers” based on each of the 100 models, using a similar genetic algorithm to the one discussed above. The artificial observers were used to generate 1,000 optimised samples each. Averaged reaction times for the combined 100,000 samples were obtained using all 100 models. The top 25 patterns that were identified for each condition and optimisation setting are illustrated in Figure 2. Figure 3 shows the mean predicted reaction times of the top 25 patterns per condition by artificial observers.

**Figure 2.**
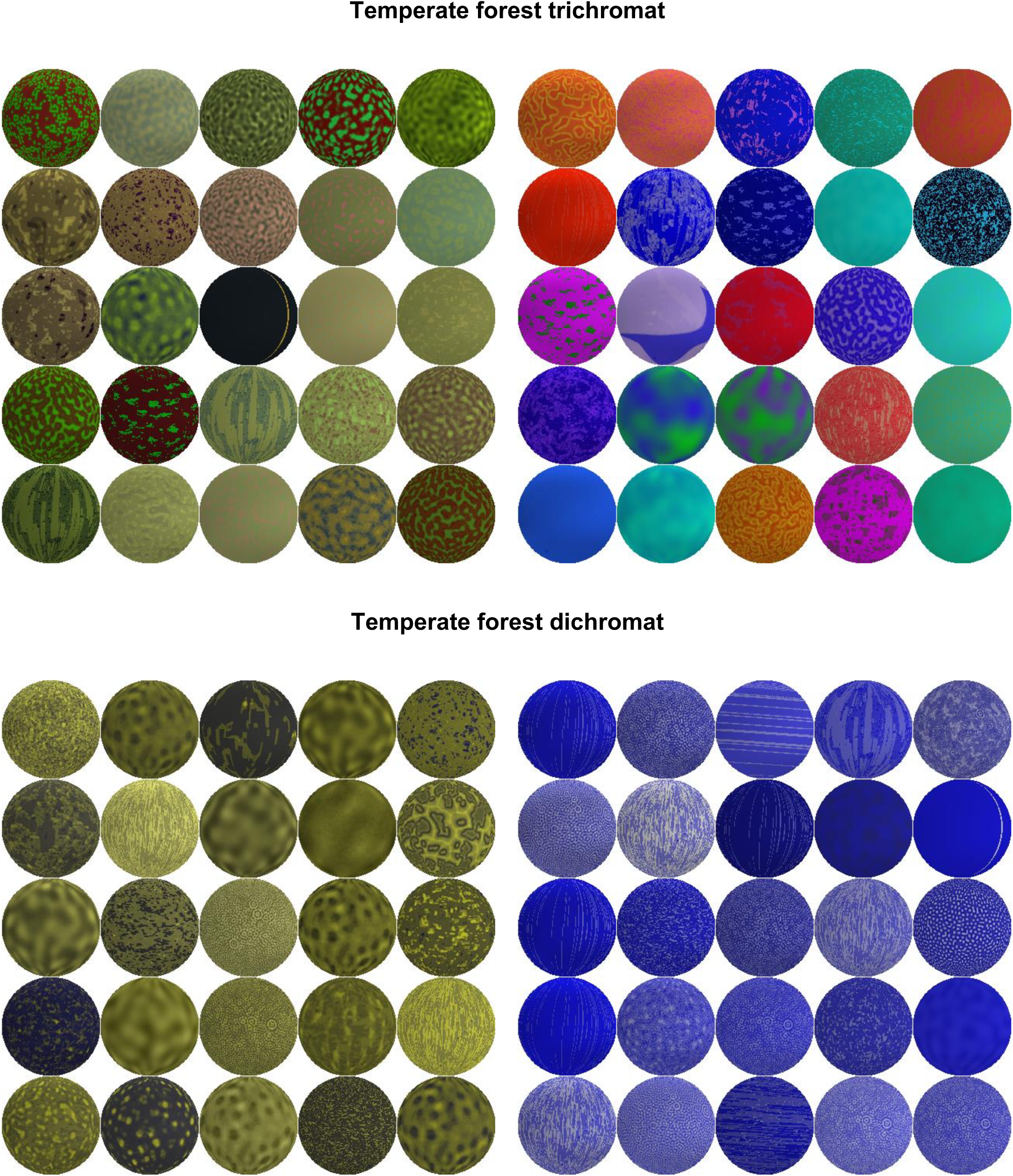
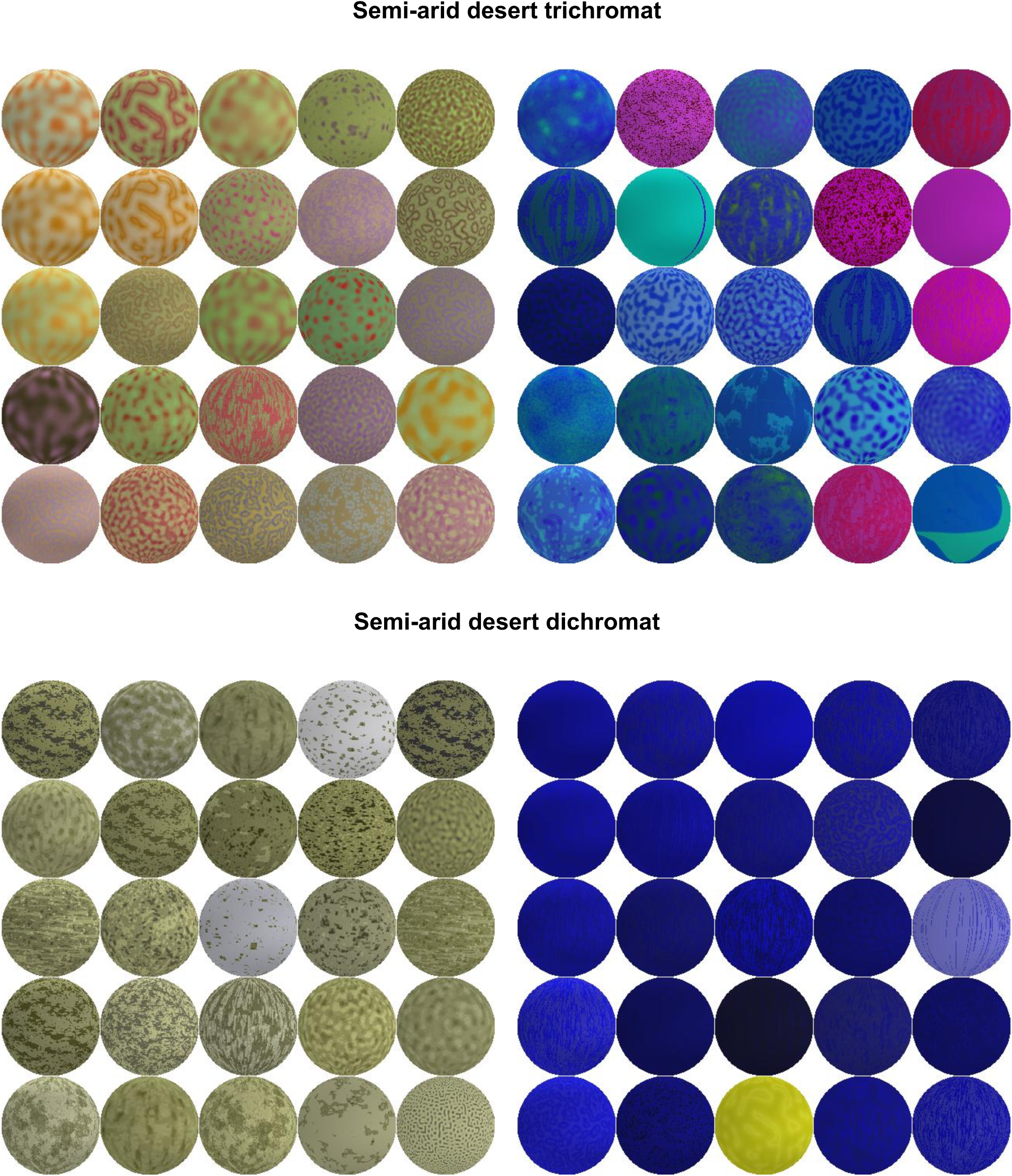
The top 25 patterns identified by the Camouflage Machine for each condition tested. Left side: hardest to see patterns; right side: easiest to see patterns. Patterns are projected on to spherical targets and shaded as shown in the experiments.

**Figure 3.**
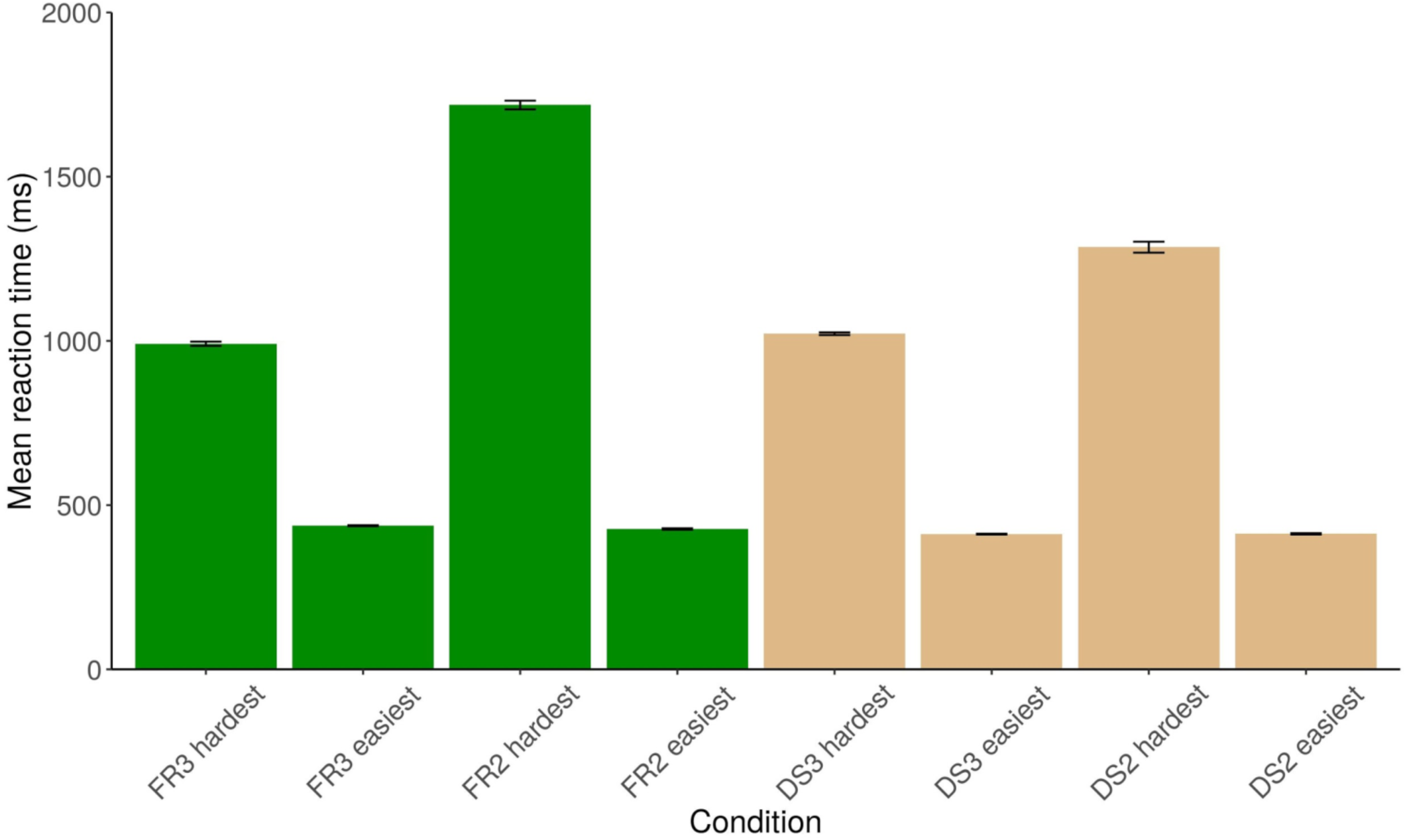
Mean predicted reaction times of the top 25 patterns per condition identified by the Camouflage Machine. Error bars are standard error of the mean. FR3: trichromatic temperate forest; FR2: dichromatic temperate forest; DS3: trichromatic semi-arid desert; DS2: dichromatic semi-arid desert.

### Output of RDNNs was validated by participants against existing camouflage

The top 25 (both optimised for hardest and easiest) patterns identified in the trichromatic temperate forest condition were tested together with two additional control patterns: British DPM camouflage and the mean colour obtained by averaging across all woodland backgrounds. DPM (Disruptive Pattern Material) was an effective camouflage pattern used by British Armed Forces for over 40 years and proven in temperate forest areas^25^. We therefore considered it an appropriate control that avoids political sensitivities created by comparisons to any current military patterns. Furthermore, the average colour of the background was khaki, which has been used by numerous militaries (including the British) from the late 19th century, making it also an important control pattern. Statistics were obtained using GLMMs, where the model, with the effect of treatment included, provided a significantly better fit than one without it (Δdeviance = 65.848, d.f. = 3, p = 3.304e-14). Post-hoc analysis (Tukey HSD) showed clearly (see Figure 4) that the hardest patterns identified by our method were significantly harder to detect than DPM (p = 0.0256) and the average colour (p = 0.0474). Similarly, the easiest patterns according to our method were significantly easier to detect than DPM (p < 0.001) and the average colour (p < 0.001).

**Figure 4.**
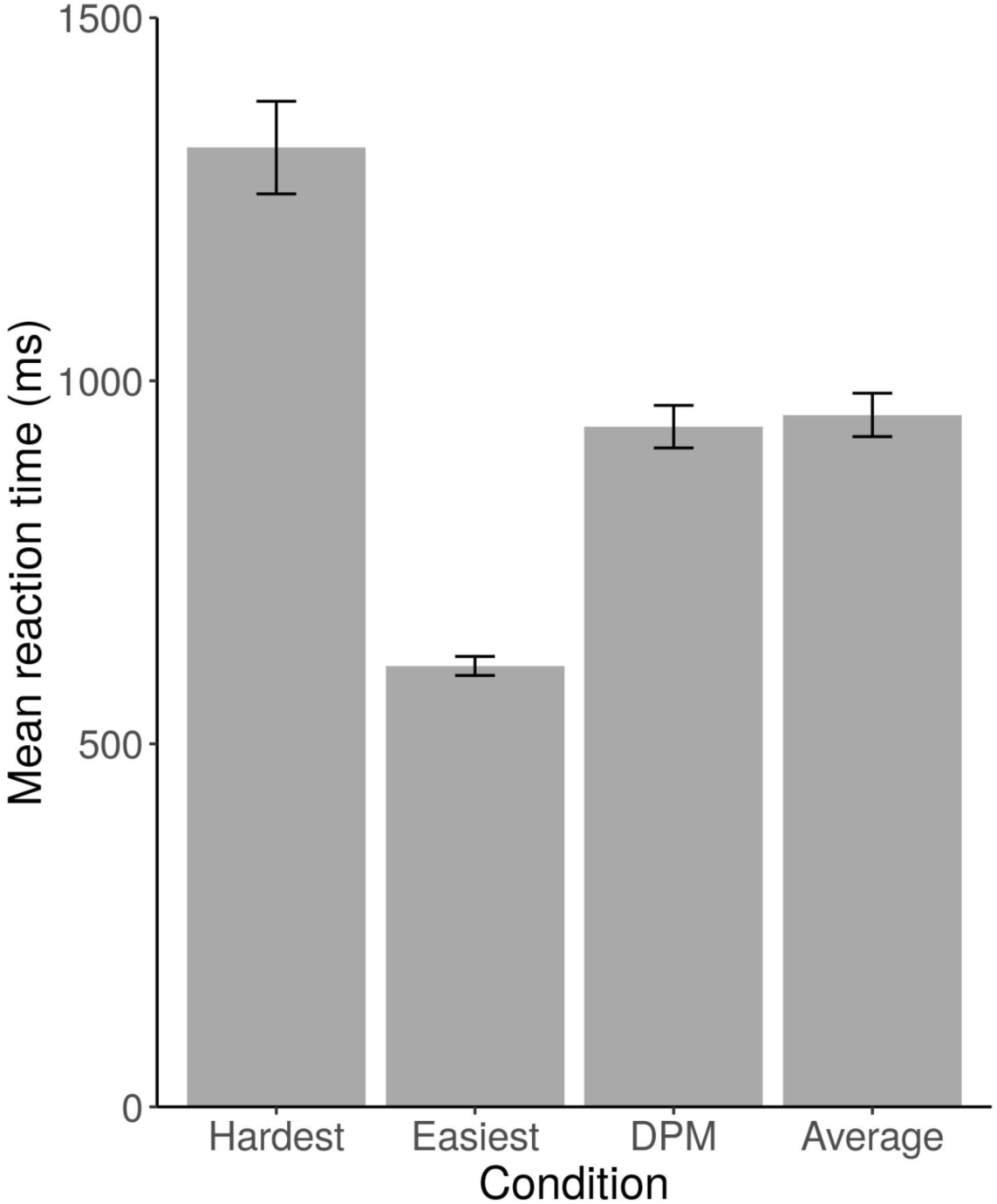
Mean reaction times to conditions tested in the validation experiment. Error bars are standard error of the mean.

## Discussion

The method presented here, named the Camouflage Machine, was successfully used to identify patterns that were better, in terms of camouflage, than an existing military camouflage pattern and the average background colour, commonly regarded as a good solution for concealment^26^. The Camouflage Machine provides an effective and efficient way to search very large parameter spaces in order to establish optimal patterns for camouflage, as well as conspicuity, in various environments. As illustrated by our simulated dichromat experiments and use of two very different backgrounds, the method generalises to different colour vision systems and across dissimilar environments. It is important to note that the Camouflage Machine need not identify a single best concealed/visible pattern, but can reveal multiple, similarly effective solutions. It is equipped to deal with the possibility that natural images contain too much variation to expect any method, including evolution, to come up with a unique, best solution. Figure 2 illustrates that there is considerable visual variability between patterns within conditions, but not in terms of predicted difficulty (see Figure 3).

In all cases, we found that the standard error of the predictions was less than 17 milliseconds, constituting what we believe to be an indistinguishable perceptual difference in the context of visually complex and non-affective stimuli^27,28^. It is also interesting to note that the predicted mean reaction times for the easiest to find patterns in each condition are equivalent. We believe this should be expected because a sufficiently salient stimulus in a complex scene should exhibit a pop-out effect^29,30,31^. Although dichromat targets optimised for concealment were significantly harder to detect than trichromat ones, consistent with our previous findings on uniformly coloured stimuli^32^, it should be stressed that our results are for trichromats using the visual information available to a dichromat, not natural dichromats neurophysiologically adapted to, and familiar with, using that level of information.

Previous studies have used evolving prey^33,34,35^; however, an important benefit of the Camouflage Machine is that far larger parameter spaces can be explored, effectively predicting data for unseen stimuli. Although deep neural networks are capable of modelling a large parameter space, establishing optima in a principled way remains a challenge. While it is technically possible to exhaustively predict every possible pattern in a given parameter space^32^, it is certainly impractical in a reasonable timescale for the space described in this study. Our solution involves combining genetic algorithms with the deep neural networks, effectively training “artificial observers”. Artificial observers allow us to be able to navigate the parameter space in a principled way and establish the hardest and easiest colour pattern combinations within reasonable timescales. For example, the predicted two-colour stimuli (optimised for concealment) were able to outperform an existing military pattern^25^ developed specifically for the (temperate forest) environment used in the experiment. We found that our genetic optimiser worked well in producing increasingly harder or easier to find patterns However, in a single condition, dichromat stimuli optimised for conspicuity in the semi-arid desert, an improvement in pattern detectability across all trials was not found. We believe the explanation for this stems from the narrower range of patterns that provide significant levels of concealment; in other words the optimiser has to deal with a space where most patterns are highly visible and so was already at ceiling performance for the majority of trials.

The Camouflage Machine offers a novel and useful tool for scientific and commercial applications. Biologists will be interested in testing various hypotheses about the colouration of animals in specific environments. For example, finding an optimal concealing pattern in an environment and comparing it to the camouflage of animals inhabiting that environment, could reveal more about their visual ecology^36,37,38,39,17,40,18^. Our method also applies to the development of bespoke camouflage for human applications, including, for example, hiding visually unappealing infrastructure^41^. Maximising conspicuity will be beneficial in safety applications, such as more salient warning signs and clothing for (motor)cyclists.

The Camouflage Machine is also capable of contributing to the development of dual-purpose applications, where both concealment and visibility is simultaneously required. An example of this is hunter’s clothing, where the wearer should be concealed from dichromatic game, but highly visible to other hunters^42^. Similarly, the Camouflage Machine can be used to further investigate distance-dependent defensive colouration^43^. Introducing viewing distance as a variable in the models would allow identification of patterns that are conspicuous close up, but become concealed at a distance^11-14^. While we deliberately limited ourselves to two colours and a simple (spherical) shape, it is clearly possible to include a larger number of colours and more complex shapes. Added to this, measures other than reaction time can be used, for example aesthetic preference^44^.

## Conclusion

The impracticality of using large arrays of patterns has previously been a limiting factor in camouflage research and studies of the adaptive value of colouration more generally^1^. We believe that our proposed method will enable larger-scale studies to be carried out. With the aid of genetic algorithms and deep neural networks, we have also demonstrated a novel approach to psychophysics, carried out using multiple dimensions. We have achieved this using a modest number of optimised samples collected from relatively few participants. Using the Camouflage Machine, it is possible to identify clusters of global optima efficiently for both concealment and conspicuity.

## Methods

### Participants

A total of 95 participants (71 female, 24 male) were recruited from members of the University of Bristol. All participants had normal or corrected-to-normal vision. Informed consent was obtained from all participants as stated in the Declaration of Helsinki. All experiments were approved by the Ethics Committee of the University of Bristol’s Faculty of Science.

### Stimuli

The creation of stimuli used the same approach as Fennell and colleagues^32^. Stimuli were created from three layers. A background layer consisted of a natural scene taken from one of two locations: Leigh Woods (North Somerset, UK, 2°38.6’ W, 51°27.8’ N) and Tabernas Desert (Almería, Spain, 2°41.3’ E, 37°02.9 N). A foreground layer was created by using a large blue cotton screen (1.8m × 2.8m) shifted across the same background. All natural images were captured with a Nikon D90 digital SLR camera (Nikon Corp., Tokyo, Japan) at a 4288×2848 pixel resolution, mounted on a tripod. The natural images captured for both background and foreground were cropped to 1920×1080 pixels prior to further processing. Between the foreground and background, a target layer was constructed from colours and textures (see below). We pre-processed the blue screen images in order to create a mask for all possible locations for the centres of targets. The derived mask allowed rapid location selection and introduction of occlusion in the foreground. A bespoke program, written using the Psychtoolbox-3 extension^45^ for Matlab^46^, was used to construct and present the stimuli, and to collect experimental data.

During all experiments, stimuli were dynamically constructed from the three layers. Backgrounds were randomly chosen from a pool of 64 images (per geographical location). Using the associated mask, a location for the target was randomly selected. Based on the number of backgrounds and potential target positions there were a very large number of potential unique stimuli.

The target was always a sphere with a radius of 64 pixels. After applying colours and texture (specific to the experiments described below), we added pseudo-realistic shading in order to produce a spherical look. The shape of a sphere was chosen as it was straightforward to create and provide it with a scene-appropriate shading. Maintaining the spherical shape throughout the experiments managed the potential confound of a target appearing different from varying angles.

Where dichromatic images were used, representations of the stimuli were created using the protan equation^47^, which simulates a trichromatic representation of an image perceived by a protanopic dichromat.

### Textures

We implemented the Gray-Scott model of reaction diffusion^48^ in CUDA^49^ C.

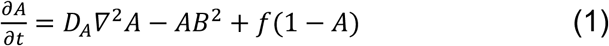

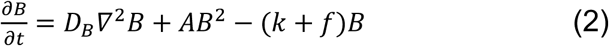

These equations calculate the concentration of two substances (A and B) based on the substances diffusion rate (D), feed rate (f) and kill rate (k), over time. The model not only represents the process of a chemical reaction but, as identified above, also produces patterns that are biologically plausible. To make the process clear, consider an initial space (image) containing random amounts of substances A and B (e.g. white noise). Over time (perhaps each iteration of a loop), one substance (A) feeds the reaction at a given rate, while the other substance (B) is killed at a given rate. Two quantities (pixels) of B can react with one of A, converting A to B (using a Laplacian Operator, ∇^2^).

Diffusion rates were fixed at 0.073 and 0.031, respectively, and every image started from the same white noise template with 10,000 iterations. The feed parameter varied between 0 and 0.25, while the kill parameter was set between 0 and 0.07. The resulting space yielded a large number of diverse textures that could be represented by a greyscale image. Many of the realised textures were homogenous and so only a subset (n = 6,809) of the textures was selected for the study (see Figure 5).

**Figure 5.**
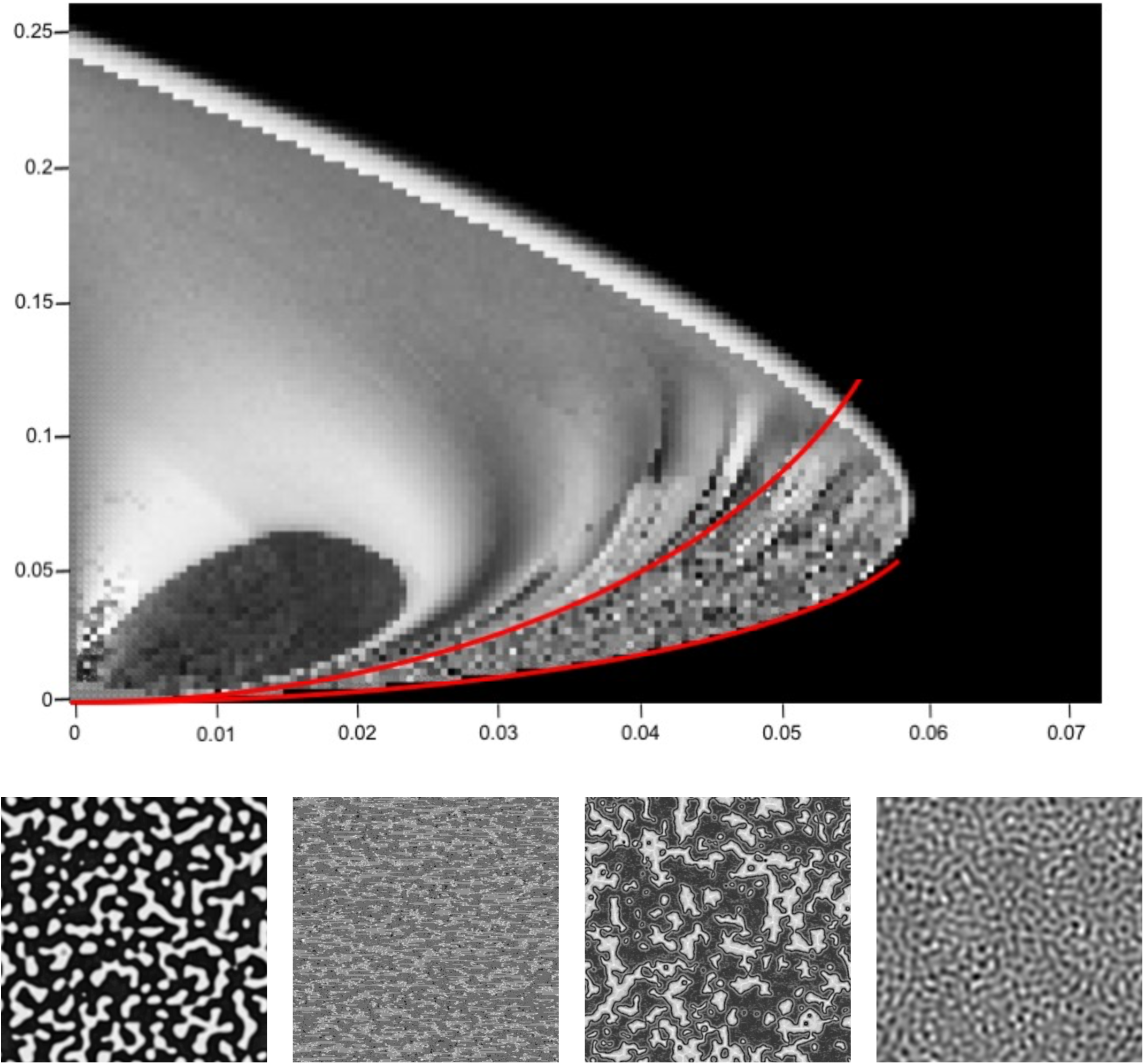
Top: The generated Gray-Scott space. To identify the region of interest, 10 points were chosen for each of an upper and lower bound for the portion of the space. The points were used to fit an exponential curve with one term i.e. of the form f(x)=a*exp(b*x) in order to obtain the a and b coefficients. The coefficients were used in a CUDA C program as limits to generate our pattern space. Unwanted patterns that were created by the CUDA program (e.g. those that are completely black) were removed. Bottom: Examples of textures used in the study.

One of the characteristics of the Gray-Scott model was that the feed and kill parameters did not provide a smooth space (i.e. adjacent patterns can be drastically different), which was necessary for optimisation. We re-parameterised the texture space to make it smoother using the following procedure. Each pattern was analysed using a Log-Gabor filter bank, providing a 24-dimensional representation of each pattern in terms of spatial frequency and orientation^50^. The resulting values were normalised and binned, in order to reduce the number of textures that could be considered perceptually very similar. This reduced the number of textures to 2,196. The 24-dimensional representation was then reduced to 3 dimensions, using t-Distributed Stochastic Neighbor Embedding (t-SNE), an effective method for dimensionality reduction popular in machine learning^51^, with perplexity of 1000 and learning rate of 1000, derived empirically. The motivation for using t-SNE rather than another popular method such as Principal Component Analysis (PCA) was that, unlike PCA, t-SNE is a probabilistic non-linear method that allowed us to create a texture space where local structure (rather than overall variance) is preserved.

### Optimisation

Parameters for the first generation of stimuli were randomly selected from the parameter spaces for each of the experiments identified below (e.g. three for each colour and three for each pattern), and the time taken to identify the stimulus recorded (fitness). A new generation was generated every fifty trials, where individual samples were selected with a genetic algorithm using tournament-based selection and tournament size of 4. Tournament-based selection is an efficient method of selecting an individual from a population of individuals in a genetic algorithm^52,53^. Tournament-based selection involves running “competitions” between members of a population, chosen at random, where the winner of each competition, the member with the best fitness, is selected for crossover. A larger tournament size reduces the probability that weak individuals will be selected (since there is a higher probability that a stronger individual is also in that tournament), thereby increasing selection pressure. Offspring, through the crossover process, received 50% of genes from each parent e.g. the best two individuals from the tournament, selected randomly. This was followed by a mutation rate of 10%, which assigned random values (mutations) to genes, randomly. The genetic algorithm was run for various numbers of generations dependent upon the experiments described below.

### General procedure

Images were projected on to a 1900×1070 mm screen (Euroscreen, Halmstad, Sweden) from 3100 mm using a 1920×1080 pixel HD (contrast ratio 300,000:1) LCD projector (PT-AE7000U; Panasonic Corp., Kadoma, Japan). For Yxy measurements of projected colours, see Table S1 in Supplementary information. Participants sat 2 m away from the display screen, so that the experimental stimulus subtended a visual angle of 50.89° by 28.59° and the target sphere 3.64°. A central fixation cross on a mid-grey background was displayed for 2 s prior to stimulus onset. Participants were asked to indicate on which side of the screen they saw the target, using the left and right shift keys on a keyboard. Each trial had a 10 s timeout; if this was reached, the experiment automatically advanced. The inter-trial interval was set to 2 s. Failure to respond was recorded as a failure and the experiment moved on the next stimulus. Reaction times were recorded to the nearest millisecond and errors indicating choice of the wrong side of the screen were logged.

### Experiments

For each experiment (unless stated otherwise), half of the participants saw targets optimised for increasing difficulty, while the other half were presented with targets optimised for increased visibility. Occlusion levels were maintained between 25 and 50% of the target, chosen randomly from a uniform distribution. Experiment 1 had 10 participants (eight females, two males) with targets of a single colour presented on temperate forest backgrounds in trichromatic colour, optimised over 500 trials. Experiment 2 had 10 participants (eight females, two males) featuring monochrome stimuli with evolving textures presented on temperate forest backgrounds, optimised over 500 trials. Experiment 3 had 10 participants (seven females, three males) who were shown targets with a fixed disruptive texture and two colours against a temperate forest background in trichromatic colour, optimised over 1000 trials. In this experiment, all participants were shown targets optimised to be hard to see.

After we confirmed that the optimiser worked, the main experiment (Experiment 4) followed a 2×2 design with two types of backgrounds (temperate forest or desert scrub) and two colour vision conditions (trichromatic or dichromatic). Forty participants (seven males, 33 females) were randomly divided between the four conditions. Each participant completed 1000 trials.

### Deep neural networks

The residual deep neural networks were written in Python 3 (Python Software Foundation, Wilmington, DE, USA) using neural network API Keras^23^. Each network was of the same configuration and consisted of an input layer, a number of residual blocks and an output layer. The input layer was 22 units, comprising: three dimensions for the pattern colour representing substance A, as described above for the Gray-Scott model; three dimensions for the pattern colour representing substance B in the Gray-Scott model; three dimensions for the texture; a dimension for level of occlusion; a two element one hot array to indicate the optimisation (hardest or easiest); and a 10 element one hot array to identify the participant. A one-hot array is a 1 × N array used to distinguish each category in a set (size N) from every other category in the set. The vector consists of zeros in all vector locations except for a single 1 in the location used to uniquely identify the category. Input colours, both trichromat and simulated dichromat, were represented as RGB triplets, with simulated dichromat values consisting of R and G channels of the same value. An alternative colour space such as CIELab or HSV could have been used, but as neural networks form their own internal representations of distances^54^ the choice of colour space is irrelevant.

Residual blocks each comprised two dense layers, a dropout layer and a summation layer, containing 768 units each, and an output layer consisting of a single variable representing difficulty as reaction time (Figure 6). We used the built in ‘rmsprop’ optimiser from Keras with the ‘mean squared error’ loss function, on difficulty, to train the networks, based on a batch size of 128 for 500 epochs.

**Figure 6.**
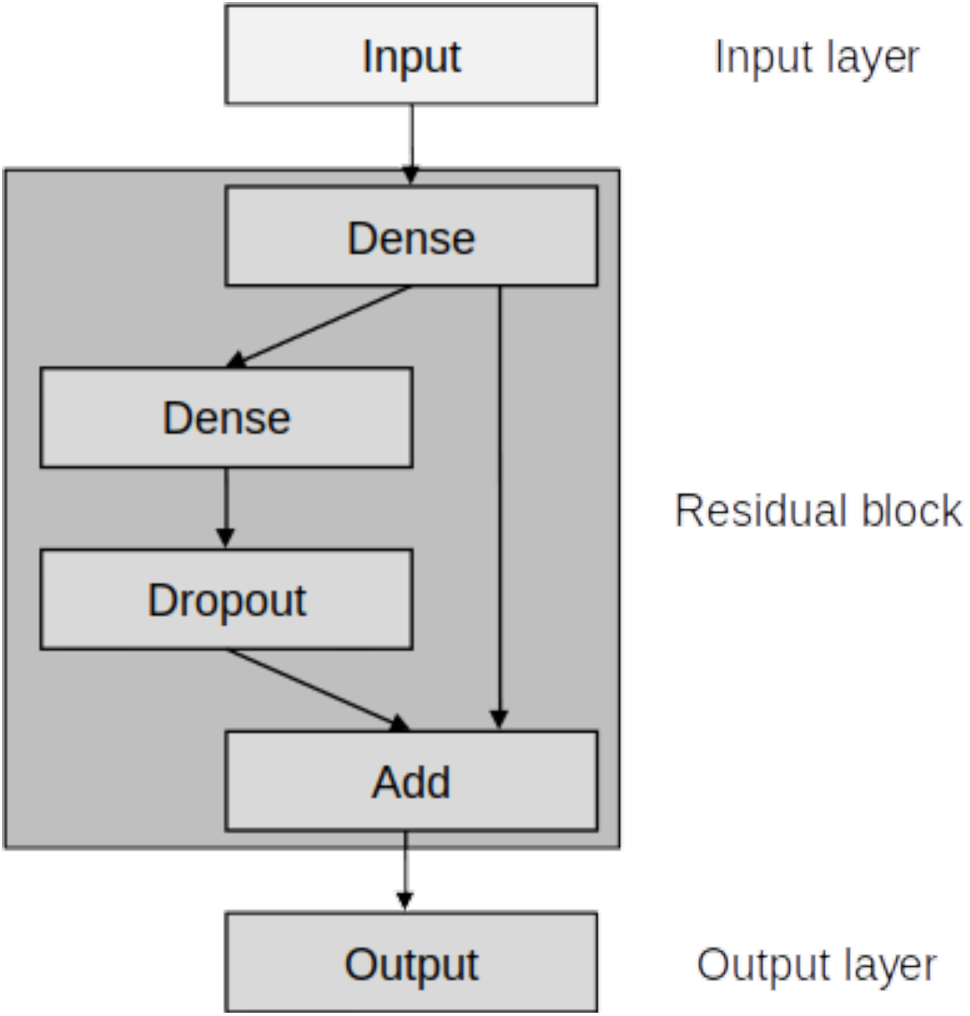
Schematic illustration of the residual deep neural network used in the study.

In order to establish the number of residual blocks to use, networks were trained with one, two, four and six residual blocks. When training a network model, a proportion of the dataset is “held-out” for validation. The training loss is the error on the training set of data, in the present case calculated using mean squared error, while the validation loss is the error, calculated in the same way, after running the held-out validation set through the trained network. As the number of epochs increases, it is expected that both the validation and training error will drop. Put simply, if validation losses are compared across different models trained with the same data, the model with the lower loss would be preferred. Here, mean validation losses were calculated for 100 bootstrapped neural networks after 500 training epochs using mean squared error (Figure 7). Statistics to compare the effects of residual block number were calculated using random permutation tests, based on 100,000 resamples. p-values were adjusted for multiple comparisons with False Discovery Rate^55^. We found that neural networks with two residual blocks produced significantly lower error rates compared to networks with one or six residual blocks, in all four experimental conditions (Table 2). While networks with two residual blocks produced significantly lower error rates compared to networks with four residual blocks in temperate forest conditions, the difference was not significant in semi-arid desert conditions. Therefore, applying Occam’s razor, we used networks with two residual blocks as it was simpler.

**Table 2.** Comparisons of mean validation losses for networks with two versus one, four and six residual blocks in all four experimental conditions.

| Condition | Comparison | p value |
| --- | --- | --- |
| Temperate forest trichromat | 2 vs. 1 | < .0001 |
|  | 2 vs. 4 | 0.0054 |
|  | 2 vs. 6 | < .0001 |
| Temperate forest dichromat | 2 vs. 1 | < .0001 |
|  | 2 vs. 4 | < .0001 |
|  | 2 vs. 6 | < .0001 |
| Semi-arid desert trichromat | 2 vs. 1 | < .0001 |
|  | 2 vs. 4 | 0.0518 |
|  | 2 vs. 6 | < .0001 |
| Semi-arid desert dichromat | 2 vs. 1 | < .0001 |
|  | 2 vs. 4 | 0.1694 |
|  | 2 vs. 6 | < .0001 |

**Figure 7.**
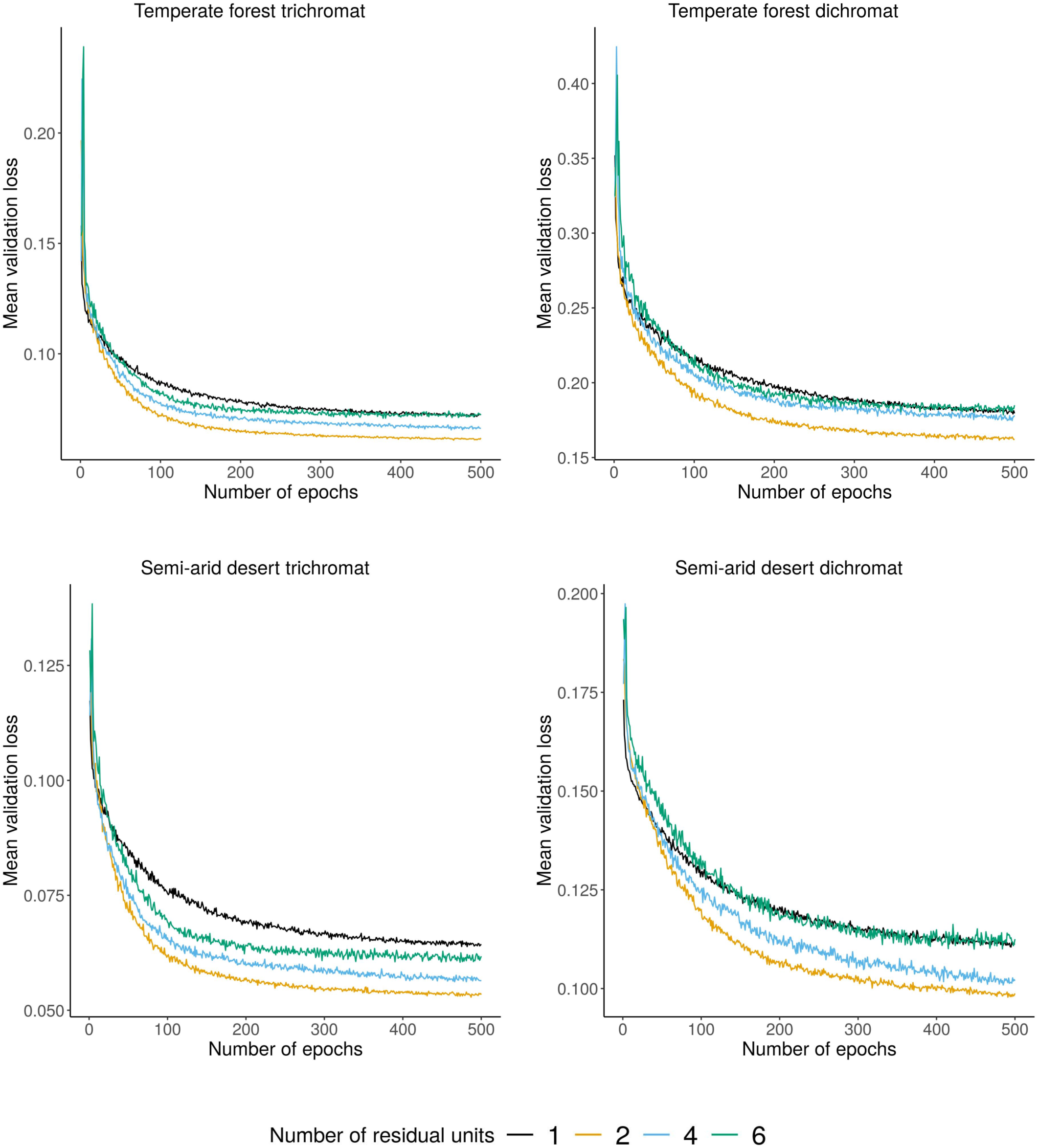
Mean validation losses for neural networks with one, two, four or six residual blocks across 500 training epochs for all four experimental conditions.

### Validation experiment

The top 25 hardest and easiest to find patterns predicted by our method from the temperate forest trichromat condition were paired with 25 DPM and 25 averaged patterns (Figure 8) for an experimental run with human participants. One run contained each pattern four times in a random order (totalling 100 trials), supplemented by four randomly selected patterns from each condition presented at the start as practice trials. We recruited 25 participants (15 female, 10 male) for the validation experiment, where each run was presented to a single participant. In all other aspects the experiment was identical to those described above.

**Figure 8.**
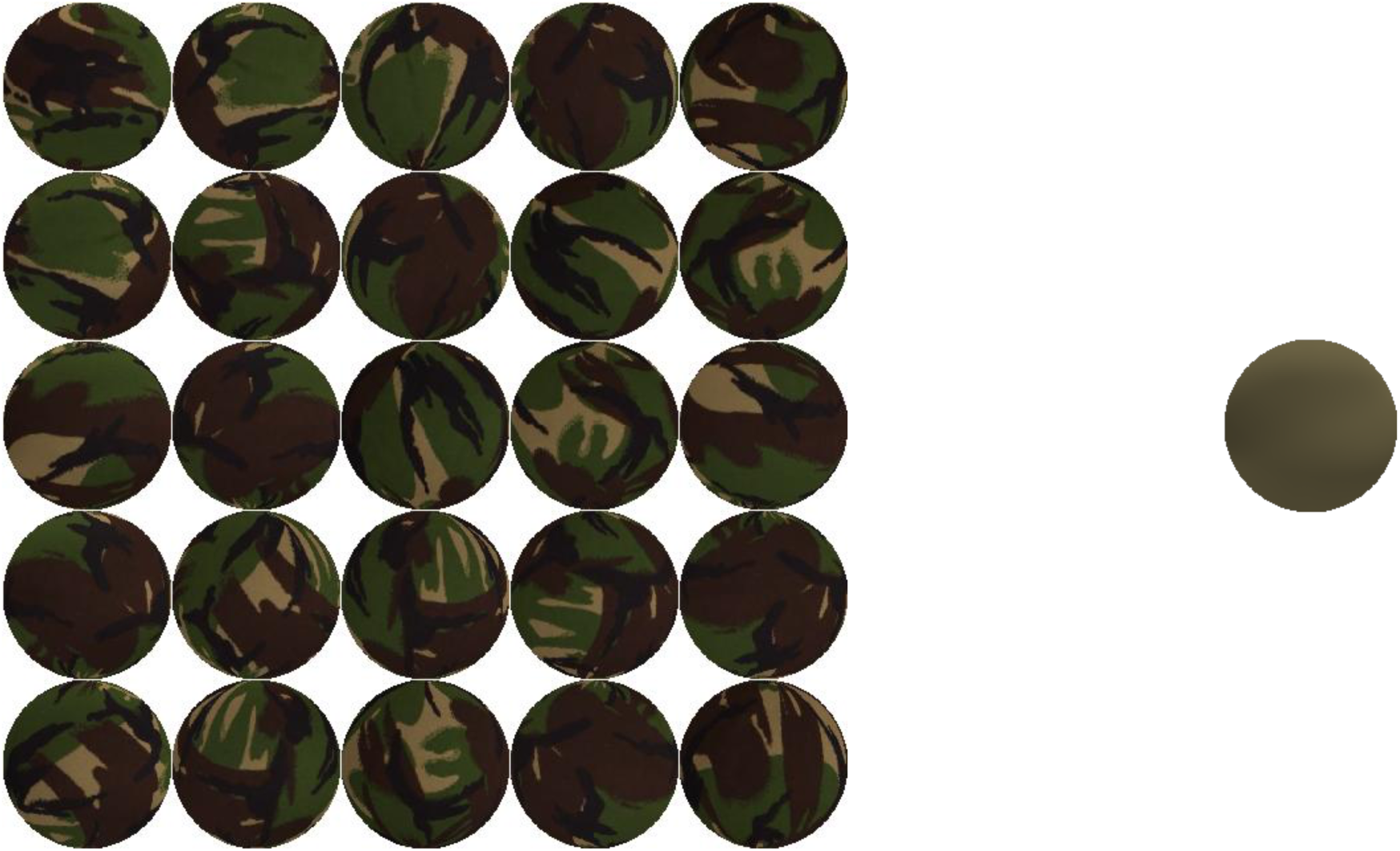
The 25 DPM patterns (left) and average colour pattern (right) presented in the validation experiment.

## Code availability

The Matlab for running experiments and Python code to train and predict from deep neural networks is available from the authors upon request.

## Supplementary Information

**Table S1.** Reference and measured values of projected colours using a Minolta CS-100A Luminance and Color Meter (Minolta Co., Ltd., Osaka, Japan).

| Manufacturer's sRGB D65 colour values |  |  | Measured Yxy values |  |  | Description |
| --- | --- | --- | --- | --- | --- | --- |
| 52 | 53 | 53 | 4 | 0.278 | 0.333 | Black |
| 84 | 86 | 87 | 10.3 | 0.279 | 0.324 | Neutral 3.5 |
| 121 | 121 | 121 | 21.8 | 0.281 | 0.326 | Neutral 5 |
| 162 | 163 | 162 | 38.4 | 0.28 | 0.33 | Neutral 6.5 |
| 203 | 204 | 203 | 60.4 | 0.276 | 0.334 | Neutral 8 |
| 249 | 249 | 244 | 78 | 0.276 | 0.343 | White |
| 0 | 137 | 167 | 22.6 | 0.2 | 0.268 | Cyan |
| 190 | 87 | 152 | 17.2 | 0.294 | 0.202 | Magenta |
| 241 | 201 | 25 | 52.6 | 0.433 | 0.537 | Yellow |
| 174 | 60 | 61 | 10.6 | 0.501 | 0.326 | Red |
| 76 | 152 | 74 | 24.2 | 0.303 | 0.552 | Green |
| 49 | 68 | 151 | 8.3 | 0.167 | 0.138 | Blue |
| 230 | 160 | 45 | 27.4 | 0.465 | 0.497 | Orange yellow |
| 162 | 190 | 65 | 34.5 | 0.38 | 0.563 | Yellow green |
| 93 | 61 | 105 | 6.2 | 0.25 | 0.201 | Purple |
| 195 | 83 | 97 | 13.2 | 0.423 | 0.296 | Moderate red |
| 72 | 92 | 174 | 11 | 0.178 | 0.159 | Purplish blue |
| 222 | 123 | 51 | 19.2 | 0.304 | 0.439 | Orange |
| 98 | 191 | 170 | 29.7 | 0.246 | 0.367 | Bluish green |
| 130 | 129 | 175 | 17.3 | 0.233 | 0.234 | Blue flower |
| 95 | 109 | 68 | 10.7 | 0.34 | 0.464 | Foliage |
| 92 | 123 | 156 | 16.2 | 0.22 | 0.253 | Blue sky |
| 195 | 147 | 129 | 23.8 | 0.356 | 0.354 | Light skin |
| 117 | 85 | 72 | 8.5 | 0.374 | 0.362 | Dark skin |

